# Deciphering the *cis-*regulatory landscape of natural yeast Transcript Leaders

**DOI:** 10.1101/2024.07.03.601937

**Authors:** Christina Akirtava, Gemma May, C. Joel McManus

## Abstract

Protein synthesis is a vital process that is highly regulated at the initiation step of translation. Eukaryotic 5’ transcript leaders (TLs) contain a variety of *cis*-regulatory features that influence translation and mRNA stability. However, the relative influences of these features in natural TLs are poorly characterized. To address this, we used massively parallel reporter assays (MPRAs) to quantify RNA levels, ribosome loading, and protein levels from 11,027 natural yeast TLs *in vivo* and systematically compared the relative impacts of their sequence features on gene expression. We found that yeast TLs influence gene expression over two orders of magnitude. While a leaky scanning model using Kozak contexts and uAUGs explained half of the variance in expression across transcript leaders, the addition of other features explained ∼70% of gene expression variation. Our analyses detected key *cis*-acting sequence features, quantified their effects in vivo, and compared their roles to motifs reported from an *in vitro* study of ribosome recruitment. In addition, our work quantitated the effects of alternative transcription start site usage on gene expression in yeast. Thus, our study provides new quantitative insights into the roles of TL cis-acting sequences in regulating gene expression.

## INTRODUCTION

Protein synthesis via mRNA translation is an essential process in gene expression. mRNA translation is divided into initiation, elongation, and termination steps. In rapidly growing cells, translation is largely limited by initiation, which occurs primarily through cap-dependent directional scanning. Specifically, the pre-initiation complex (PIC), which consists of the 40S ribosomal subunit and multiple initiation factors, must scan the 5’ transcript leader (5’ TL) to locate the main coding region (CDS) (1). As such, sequence elements within 5’ TLs are important contributors to initiation efficiency and gene expression. Previous work demonstrated that features such as upstream open reading frames (uORFs), mRNA structures, and mRNA binding protein motifs influence the identification of the main start codon by the PIC (1–6). These features have been studied largely independently, such that their relative importance in regulating gene expression from native 5’ TLs has not been systematically evaluated *in vivo*.

The Kozak context, a consensus sequence directly upstream of the start codon, was first identified in early studies as a key factor in translation initiation (7). When a PIC encounters a start codon in optimal Kozak context, initiation factors reorganize into a “closed” formation to initiate translation (8, 9). In contrast, PICs presented with start codons in a sub-optimal Kozak context are more likely to remain in an “open” scanning conformation and skip initiation. The process by which the PIC bypasses an upstream start codon to initiate translation downstream is described as the leaky scanning model (LSM). Previously, a massively parallel reporter assay (MPRA) measuring translation of 2,041 TL variants of the yeast RPL8A gene found that variation in the Kozak context resulted in a ∼7-fold difference in protein expression, with a preference for A-rich sequences (10). Additional experiments confirmed this and highlighted the positive influence of adenosine at the -3 position (11–14). Other work has shown translation can also initiate at near-AUG codons (15–22), though this appears to be generally inefficient in yeast (13, 23–27). Though the yeast Kozak motif itself is well defined, the relative importance of Kozak sequences in the context of other elements in native yeast TLs *in vivo* remains unstudied. For example, other studies have found mRNA secondary structure can prevent start codon recognition (28, 29). Thus, the extent to which leaky scanning and the Kozak context alone control translation from natural TLs remains unknown.

Alternative transcription start sites add another layer of complexity to the function of mRNA transcript leaders. Several studies have shown that many genes initiate transcription at multiple sites (5, 30, 31). *In vitro* measurements of 96 native yeast TLs using luciferase assays, revealed that alternative transcript initiation sites that differed by 50 to 200 nucleotides in 5’ ends could influence translation efficiency up to 100-fold (32). Large changes in expression of alternative TLs suggest that sequence features play a crucial role in this regulation. For example, alternative transcription start sites could alter the number and identify of uORFs, RNA structures, and RNA Binding Protein sites in 5’ UTRs. However, the effect of such differences in natural 5’ UTRs on protein expression has not been systematically evaluated.

While several foundational studies analyzed gene expression from 5’ TLs, they did not evaluate the relative impacts of sequence features in full-length native TLs *in vivo*. Studies using libraries containing a fixed sequence at the 5’ end may not capture the effects of structure or sequence preference near the cap. The fixed length, synthetic, and randomized 5’ TLs previously tested *in vivo* (*10, 11, 14*) were relatively short, and do not explain larger length differences or naturally occurring motifs present in native TLs. A recent study compared the recruitment of yeast ribosomes to native TLs *in vitro*, and reported several motifs associated with high and low ribosome recruitment (12). However, although useful, translation extracts do not fully reproduce the cellular environment. Furthermore, this approach required the removal of all upstream AUGs (uORFs), which are large contributors to gene regulation by native TLs. A systematic study *of in vivo* regulation by natural TLs is needed to more fully evaluate 5’ TL impacts on gene expression.

Here, we assayed protein expression, mRNA levels, and ribosome loading from a comprehensive library of endogenous yeast TLs and determined the relative impacts of TL features. We first compared expression levels from hundreds of genes containing alternative TLs and found that even slight changes in transcript leader length greatly impacted expression. Next, we evaluated the extent to which Kozak sequence strengths explain gene expression variance from *S. cerevisiae* TLs. Using start codon-Kozak pair strengths, we built a leaky scanning model which more accurately explained ribosome scanning and expression. Finally, we defined and calculated strengths of additional *cis*- and *trans*-regulatory sequence elements shown to influence expression. By combining all features in a computational model, we explain ∼70% of the variance in expression across native transcript leaders. By determining the relative impacts of TL features on gene expression, our findings shed new light on the “rules” of directional scanning.

## MATERIALS AND METHODS

### FACS-uORF and PoLib-seq data

All raw sequencing data were from NCBI SRA accession number PRJNA721222. The data analyzed in this study included wildtype UTRs containing uORFs (reported previously in (34)) and additional wildtype UTRs that do not host uORFs. Data were processed as previously described (34). Briefly, read pairs were merged and error corrected using FLASH2 (94) using parameters ‘-z -O -t 1 M 150’. The resulting merged reads were trimmed to remove the ENO2 promoter sequence (plasmid libraries, AGTTTCTTTCATAACACCAAGC) and the cDNA adapter sequence (RNA libraries, AGTTTCTTTCATAACACCAAGCNNNN) using cutadapt with parameters ‘--trimmed-only -e -0.04. RNA-seq libraries were further processed to remove extra nucleotides incorporated at the 5’ cap by reverse transcriptase (NNG, (34)). Trimmed reads were counted for perfect matches to designed library constructs using custom perl scripts (DNA-seqcount.pl and RNA-seqcount.pl; (34)). Relative YFP levels were calculated for each wildtype yeast UTR by comparison to YFP/mCherry TECAN luminometer readings taken from each of the eight FACS-sorted bins after growing the sorted yeast cultures overnight in YPD at 30°C. Each biological replicate was normalized to a 0-1 scale of YFP expression, and read counts were scaled to the proportion of cells sorted into each bin. The average YFP value for each transcript leader was calculated as follows: YFP / mCherry = (SUM(YFPbin * reads / bin), where YFPbin represents the YFP / mCherry ratio measured by the TECAN, normalized to a 0-1 scale. The YFP / mCherry levels were compared across the three replicates to remove noisy TLs with inconsistent measurements (standard deviation > 0.05, less than 50 normalized reads).

PoLib-seq estimates of ribosome loading were also calculated as previously reported (34). To summarize, reads were pooled into “translating” (disome and larger) and “non-translating” (40S, 60S, and monosome) fractions separately for each replicate and replicate 1 was downsampled by a factor of 0.826 to ensure similar proportions of translating and non-translating reads in both replicates. A 5000 total read cutoff (combining the two replicates) was used to calculate the % translating metric for PoLib- seq library measurements.

### Leaky Scanning Model (LSM)

Kozak scores ranging from 0-1 for each AUG codon were taken from our recent work (13). The leaky scanning model (LSM) is used to calculate an adjusted Kozak score for each TL. The model includes all AUGs present in the 5’ TL. uAUGs detract ribosomes from the main start codon and decrease the overall Kozak score. To calculate adjusted Kozak scores that contribute to YFP, we first calculate the probability that a ribosome has skipped all start codons upstream of a productive CDS start codon:

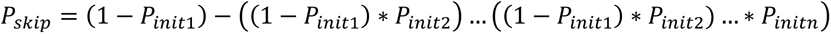

where P_init1_ representing the Kozak score of the first uORF AUG the PIC reaches in the 5’ TL (nearest AUG from 5’ end). P_skip_ represents the amount of “available/remaining” PICs that bypass upstream AUGs and continue scanning the 5’ TL. The adjusted Kozak score for initiation at an AUG that produce YFP is calculated as:

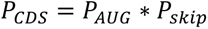

A few TLs contained unannotated N-terminal extensions due to uAUGs that are in frame, without corresponding stop codons. Because such start codons would produce functional YFP, the adjusted Kozak score for such TLs was calculated by summing P_CDS_ scores for each productive AUG. After all AUGs have been accounted for, in-frame AUG Kozak scores added to the adjusted score, while out-of-frame AUG Kozak scores are subtracted from the score. Thus, the LSM captures the leaky scanning events that ribosomes encounter when nearing AUGs.

### TL feature compilation

Kozak scores were taken from our previous reporter studies for AUG start codons (-4 to +1) representing NNNNATGN (34). Nucleotide frequencies of (A/T/C/G) were calculated and represented as a fraction of the whole TL. There were two separate frequency calculations. Cap proximal frequencies (∼20 nucleotides from 5’ end) and Distal frequencies (∼30 nucleotides near start codon). Mean RNA levels were determined by averaging data over three replicates for each 5’ TL. MaxAStretch denotes the longest stretch of consecutive A’s in the TL. G quartets (Num4Gs) was the number of times “GGGG” is found in the TL (note: GGGGG counts as 1 g-quartet). The ViennaRNA (44) package was used for computationally predicting structure around the 5’ cap and the start codon. ΔΔG of the start codon was used for predicted structure around the mainORF. This included 30 nucleotides around the main AUG. The energies of unwinding the RNA around uORF start codons (ΔΔG) were estimated based on previous work (28) using programs from ViennaRNA (44). For ΔΔG predictions, we used RNAsubopt to predict 100 suboptimal folded structures for each 5’ TL, including some of the YFP (50 nucleotides). Start codon unfolded structures were set by unpairing the -15 to +15 region around each start codon. The ΔG of each TL folded and unfolded structure was then predicted via RNAeval. The un-structured estimates were then subtracted from the structured predictions to calculate one hundred estimates of the ΔΔG for each TL. The mean values from these calculations were used as the final modeling features. The ΔG of the 5’ cap was predicted by averaging 100 trials of structure prediction via RNAfold. Motifs for over- predictors and under-predictors were identified via STREME (48) using parameters (DNA to RNA, 4-12 motif length, p-value 0.05, center align). G-quadruplexes were estimated using the QGRS package (46). Motifs identified in Hogan et al (35) were incorporated as sum of counts into the model, but were not significant in modeling.

### 5’ TL Elastic Net Regression Modeling and Feature Selection

Elastic net modeling is a linear regression-based algorithm that is a hybrid of lasso (L1) and ridge (L2) models. L1 regularization tends to limit grouped feature selection and minimizes overall significant features. On the oSr hand, L2 regularization shrinks feature coefficients, but does not force them to be zero. As such, elastic net regression overcomes limitations of selected variables and shrinkage by incorporating both lasso and ridge. The objective function minimized is:

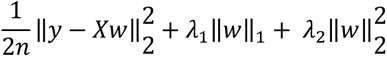

The orange and purple part correspond to L1 and L2 penalties respectively. The model was built using the ElasticNet (ElasticNetCV ) package from sklearn (95) with n=100. The function takes in two parameters (α, l1) which represents the amount of regularization penalties and the scaling between L1 and L2 penalties, respectively. The data were split into training and testing sets (68% and 32%) to avoid over-fitting. The model performance was quantified by the R^2^ score which compares the predicted versus measured YFP/mCherry levels for 5’ TLs. In total three separate EN models were created 1) Less Stringent EN Model for all 5’ TLs 2) More Stringent EN Model for all 5’ TLs (FIG S4) 3) EN Model for all 5’ TLs lacking uAUGs (FIG S5). All model data are in the supplementary table (TABLE S4,S5,S6).

## RESULTS

### Using multiple MPRAs to evaluate the effects of endogenous yeast transcript leaders on gene expression

We developed a version of FACS-seq, an MPRA system to accurately evaluate protein levels, for endogenous yeast 5’ TLs. Previously, we identified significant transcription start sites (TSSs) genome wide and re-annotated 5’ TLs in five yeast species (33). We constructed a library of 15,152 5’ TLs from *S. cerevisiae* and *S. paradoxus* upstream of YFP, on plasmids that also express mCherry as an internal control (FIG 1A). TLs that were up to 180 nucleotides long and represented at least 10% of their host genes’ transcripts were included. This includes 86% of yeast transcript leaders, enabling us to capture a wide range of gene expression and covering numerous sequence features. Yeast transformed with the transcript leader library were FACS-sorted into expression bins based on the YFP / mCherry ratio (FIG 1B). By sequencing the constructs from each bin, we calculated the mean YFP expression level for individual TLs. We performed FACS-seq in triplicate, and replicates were highly correlated (FIG S1; R^2^∼0.98-0.99; TABLE S1). Overall, the range of YFP expression among transcripts in our 5’ TL library varied over a 100-fold (FIG 1C). To our knowledge these results define, for the first time, the range of gene expression driven by natural yeast transcript leaders *in vivo*.

**Figure 1.**
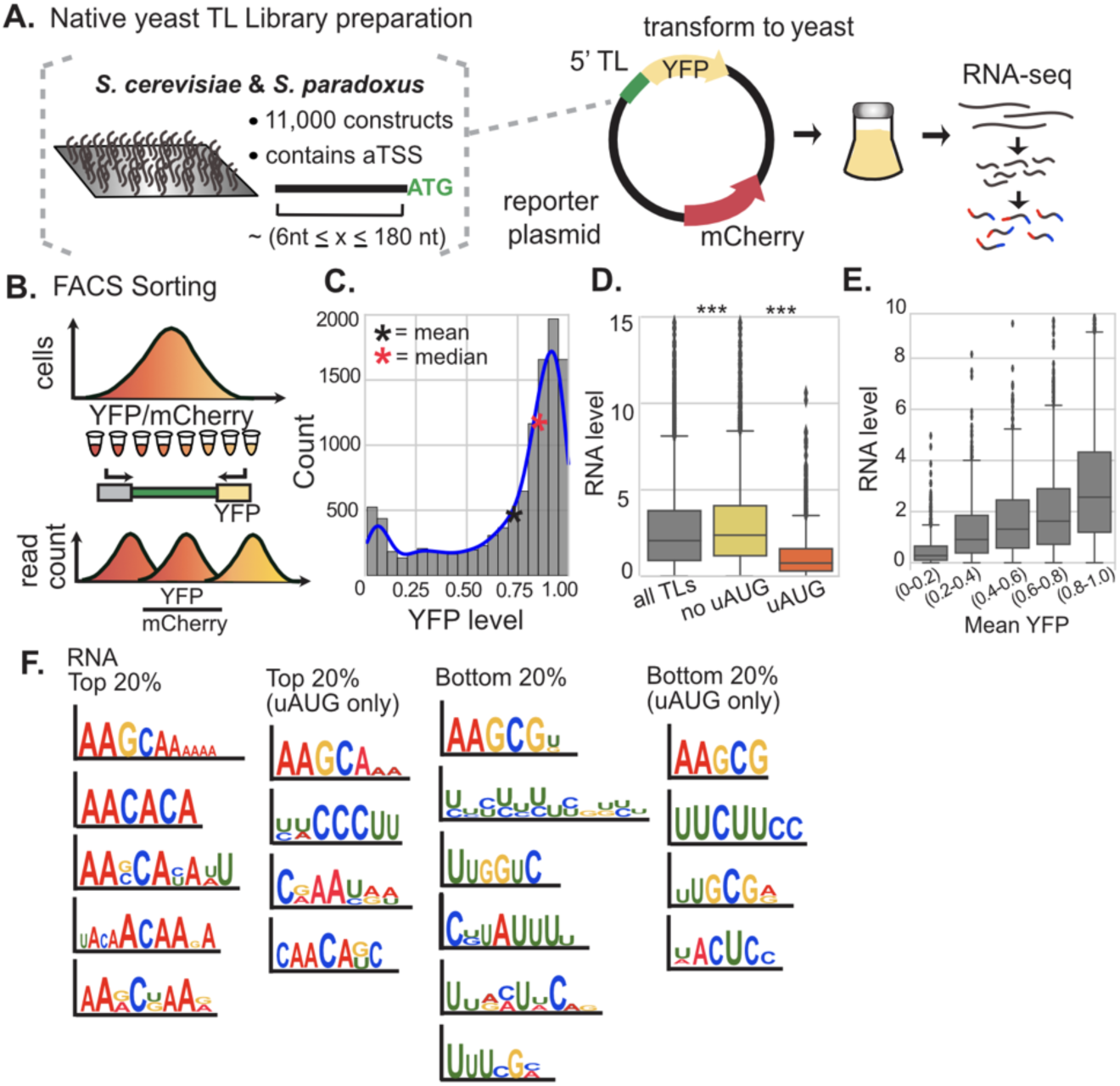
MPRA analysis of transcript leader influence on protein and RNA levels. (A) FACS-Seq - a library of thousands of native yeast 5’ TLs, including alternative transcription start sites was cloned upstream of a single-copy YFP dual fluorescence reporter plasmid containing mCherry as an internal control. The reporter plasmids were transformed into S. cerevisiae. Targeted RNA-seq was used to assay RNA levels, relative to plasmid levels (RNA_rpkm_/DNA_rpkm_) (B) Cells were sorted and binned via FACS. Plasmids were extracted and sequenced to assay YFP expression levels for each reporter. (C) The YFP distribution for the 5’ TL library (mean=0.713, median=0.842). (D) Average RNA level across 3 replicates of transcripts for different 5’ TL groups (all, no uAUG, uAUG). n=9382, 7923, 1459 (E) RNA levels for different YFP level groups. Outlier RNA-levels above 10 not shown. n = 1255, 745, 765, 1769, 6477. (F) STREME motifs identified in the Top and Bottom 20% of RNA levels with and without uAUGs. (P-values) *** < 0.001, ** < 0.01, * < 0.05

Transcript leaders can also impact mRNA levels which could contribute to variation in YFP expression. For example, yeast upstream ORFs (uORFs) induce Nonsense Mediated Decay to various extents (34). Other TL sequences may affect gene expression by recruiting RNA binding proteins (35, 36) or even causing premature transcription termination (37). To investigate this, we next performed targeted RNA-seq to estimate reporter RNA levels, normalized to plasmid DNA. Notably, this confirmed that the ENO2 promoter driving YFP used the designed transcription start sites (TSSs) 97% of the time on average (34). RNA-seq showed mean RNA levels varied ∼100-fold (FIG 1D), and positively correlated with YFP expression (R=0.392, FIG 1E). This is attributable in part to upstream ORFs (uORFs), which can induce nonsense-mediated decay (NMD) of mRNA (FIG 1D). Excluding 5’ TLs with uORFs, we found AC-rich and U-rich motifs significantly enriched in high and low mRNA levels, respectively (FIG 1F). Thus, natural yeast 5’ TLs harbor *cis-*regulatory sequence features that correlate with variation in mRNA expression levels, potentially due to effects on RNA stability.

Next, we assayed ribosome loading for TLs in our library using PoLib-Seq (34, 38). We fractionated polysomes on a sucrose gradient and calculated relative ribosome load (RRL; %translated/total; see methods) for each TL (FIG 2; FIG S2; TABLE S2). RRL varied ∼3-fold for natural yeast TLs (FIG 2B). There was a strong positive correlation between the FACS-seq and Polib-seq assays (FIG 2C; R^2^ = 0.736). This indicates that our FACS-seq YFP measurements were largely consistent with ribosome loading. In addition, we observed a complex relationship between ribosome load and mRNA levels. TLs with the lowest ribosome load also had the lowest RNA levels, mid-range RRL values (40-60%) had the highest RNA levels, and a decline in RNA abundance was observed for TLs with higher ribosome loads (FIG 2D). This was independent of the presence of uORFs (FIG 2E). This observation suggests an optimal level of ribosome loading, wherein translation appears to protect RNA from degradation up to a certain threshold, above which excessive ribosome loading may lead to RNA decay, perhaps due to ribosome collisions (39, 40). Finally, we identified sequence motifs enriched in TLs with high and low-ribosome loading (FIG 2F). In general, we found A-rich sequences, including “AUA”, were associated with high ribosome loading, while U/C-rich sequences were associated with low ribosome loading.

**Figure 2.**
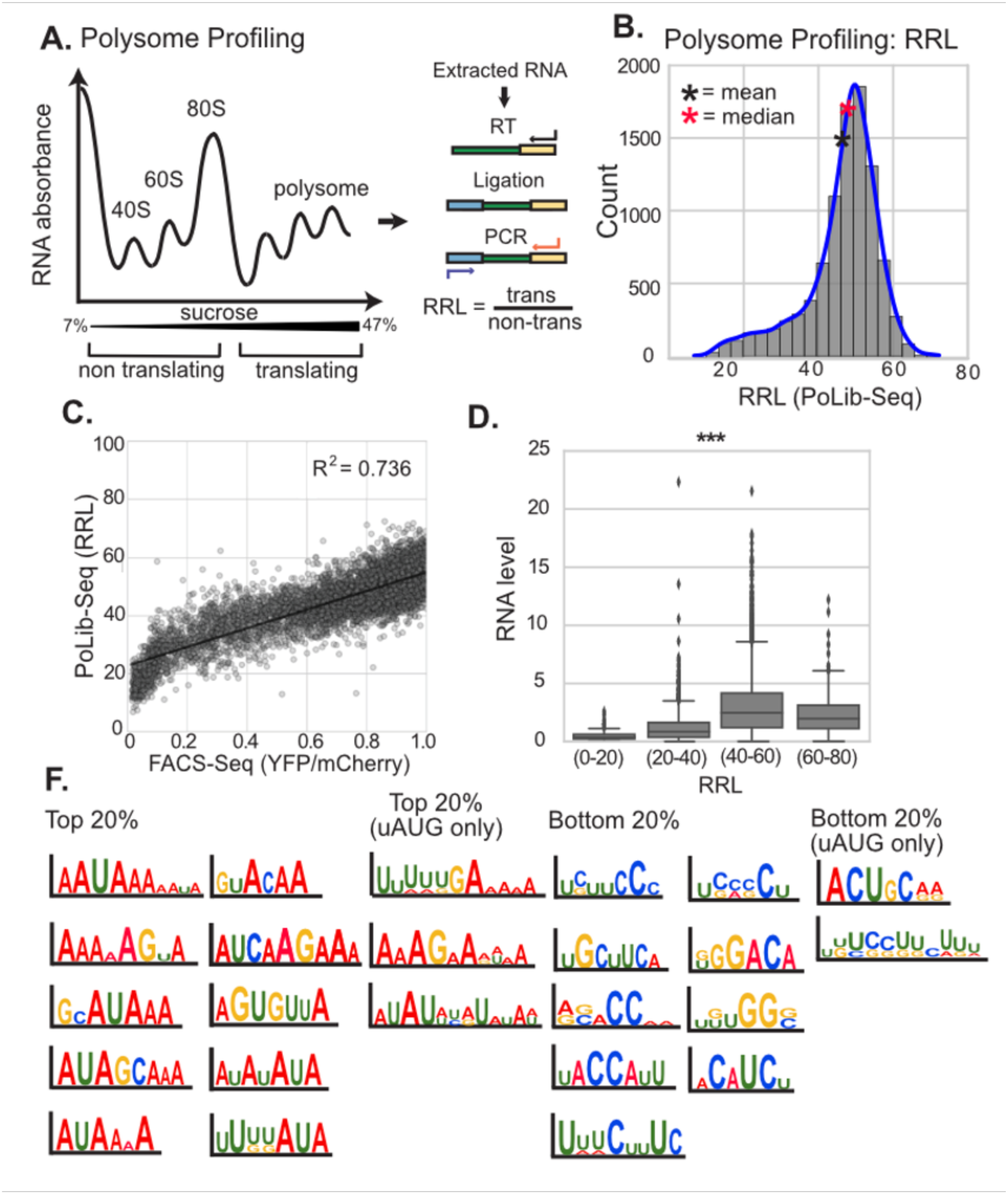
PoLib-Seq MPRA determines relative ribosome loading driven by natural yeast transcript leaders. (A) The schematic depicts the PoLib-Seq assay used to measure the ribosome loading on 5’ TLs. Polysome extracts were prepared from WT yeast strain (BY4741). The extracts were fractionated on a 7% to 47% sucrose gradient via ultracentrifugation. UV absorbance graph represents polysome fractions from non-translating (40S, 60S, 80S) to translating (2 polysomes +). RNA extracted from Polysome fractions was prepared for sequencing (methods). Relative ribosome load (RRL) was calculated as translated/total for each TL. (B) Distribution of PoLib-Seq measurements of RRL for 5’ TLs. (C) FACS-Seq vs PoLib-Seq: Comparison of YFP/mCherry (x-axis) from FACS-Seq and RRL from PoLib-Seq (y-axis) results for the 5’ TL library. The measurements are highly correlated with R^2^ of 0.736. (D) Mean RNA levels for different RRL level groups for all TLs. n = 234, 1396, 7329, 423. All p-values < 0.001 F) STREME motifs identified in the Top and Bottom 20% of RRL levels with and without uAUGs. (P-values) *** < 0.001, ** < 0.01, * < 0.05

### Evaluating the impact of 5’ TL RNA structures on YFP expression

Prior work found mRNA structures can hinder PIC scanning efficiency and limit initiation(2–4, 41–43). We next assessed how native yeast structural features affect YFP reporter expression *in vivo*. First, we used RNAfold to calculate the predicted stabilities of the TL cap (first 40 nucleotides) and start codon (+/- 15 nucleotides) regions (methods) (44)(FIG 3A & 3B). These estimated cap structural stabilities varied over more than a 100-fold range, with the most structured caps reaching -14 kcals/mol.

**Figure 3.**
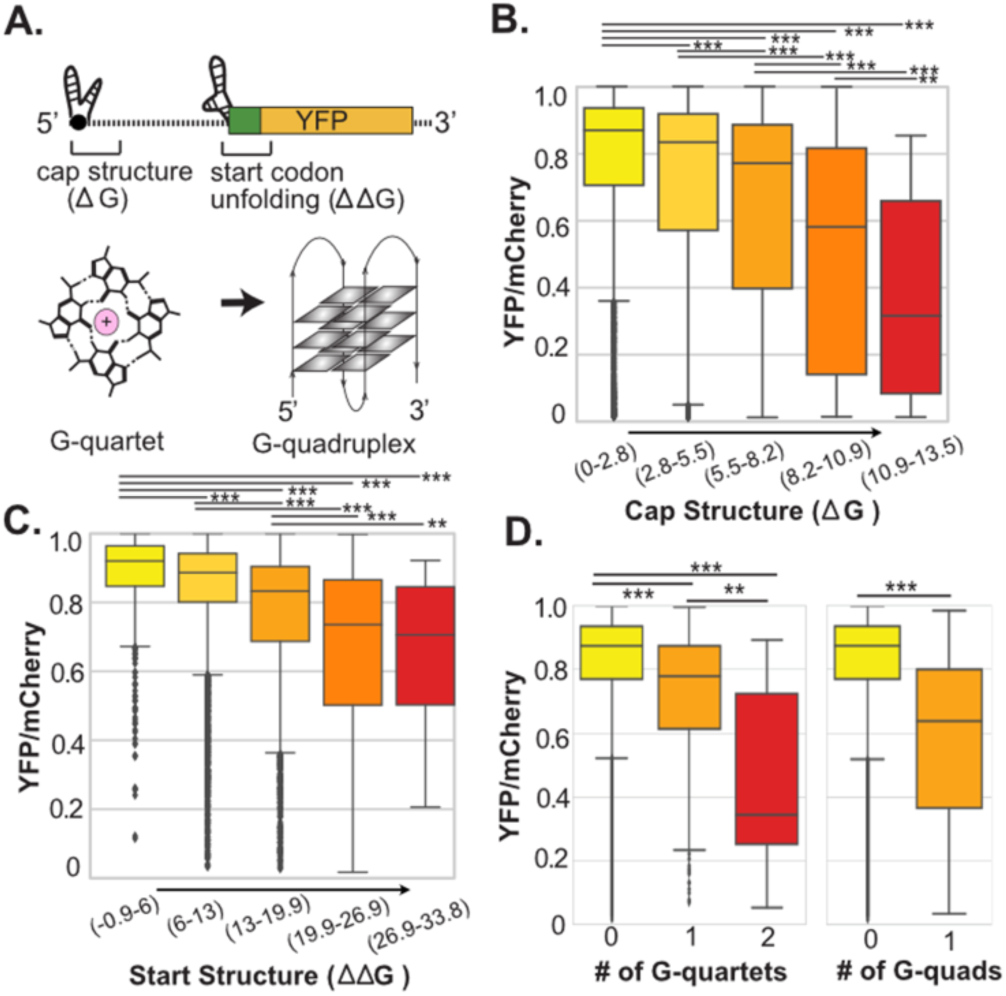
Impacts of natural yeast transcript leader RNA structure on gene expression in vivo. (A) Schematic displaying different mRNA structures: cap structure, start codon structure, g-quartets, g-quadruplexes. (B) Boxplots showing the relationship between cap structure (ΔG) and YFP levels. n = 4279, 3091, 1338, 248, 32 (no uAUG TLs included) (C) Boxplots representing the association between YFP levels and the unfolding energy of structures surrounding the main start codon (ΔΔG). n = 498, 5832, 2368, 266, 24 (no uAUG TLs included) (D) (left) G-quartet structures form when 4 Guanines (in a row) are Hydrogen bonded to each other. From the TL dataset, we saw G-quartets lead to decreased initiation. n = 8753, 292, 13 (no uAUG TLs included) (right) In the presence of metal ions, these G-quarters can stack on top of each other to form higher-order structures known as g-quadruplexes. The boxplots show a decrease in YFP expression in the presence of g-quadruplexes n= 8950, 107 (no uAUG TLs included). Only 1 gene (YBR196C-A) was predicted to contain 2 g-quadruplexes (not shown). (A-C) (P-values) *** < 0.001, ** < 0.01, * < 0.05

Structures around the start codon were estimated as the change in free energy ( (Δ-Δ-G; ΔΔG) to represent the unfolding energy required for the PIC to access the start codon (maximum =∼34 kcal/mol). Consistent with current models, we found both structured cap and start codon regions correlated with decreased YFP levels in our library (FIG 3B, 3C). Indeed, the structural stability of the TL cap region was much more repressive than stability around the start codon, suggesting structures that inhibit initial PIC loading have a larger effect on gene expression than structures that affect PIC scanning.

RNA also folds into higher-order structures such as g-quartets which can subsequently fold into stable g-quadruplexes, hindering translation (FIG 3A) (11, 45). We examined TLs with the potential to form these higher-order structures to quantify their impacts on YFP expression. TL sequences containing the GGGG motif were considered to have g-quartets. Strikingly, the addition of a single g-quartet significantly decreased YFP expression in natural yeast TLs (FIG 3D). Using QGRS Mapper (46), we predicted the formation of g-quadruplexes amongst the 5’ TLs. While g-quadruplexes were rare (<2% of TLs) in our library, their presence was associated with significantly reduced YFP expression (FIG 3D).

These results show that RNA structures in natural yeast TLs generally reduce protein expression, and show a wide range of effects associated with predicted structural stability of TL cap regions, start codon stability, g-quartets and g-quadruplexes.

### Predicting 5’ TL YFP expression using a Leaky Scanning Model

Our next aim was to quantitatively evaluate the role of Kozak sequences (7, 8) in regulating PIC scanning and initiation in natural yeast TLs. We recently defined yeast Kozak strengths from the -4 to +1 position for AUG and near-AUG start codons using FACS-Seq (13). Using these data, the Kozak strength of the main start codon alone (main codon model, FCM, FIG 4A) explained roughly 13% and 22% of expression for the endogenous yeast 5’ TLs with and without uORFs, respectively (FIG 4B; FIG S3).

**Figure 4.**
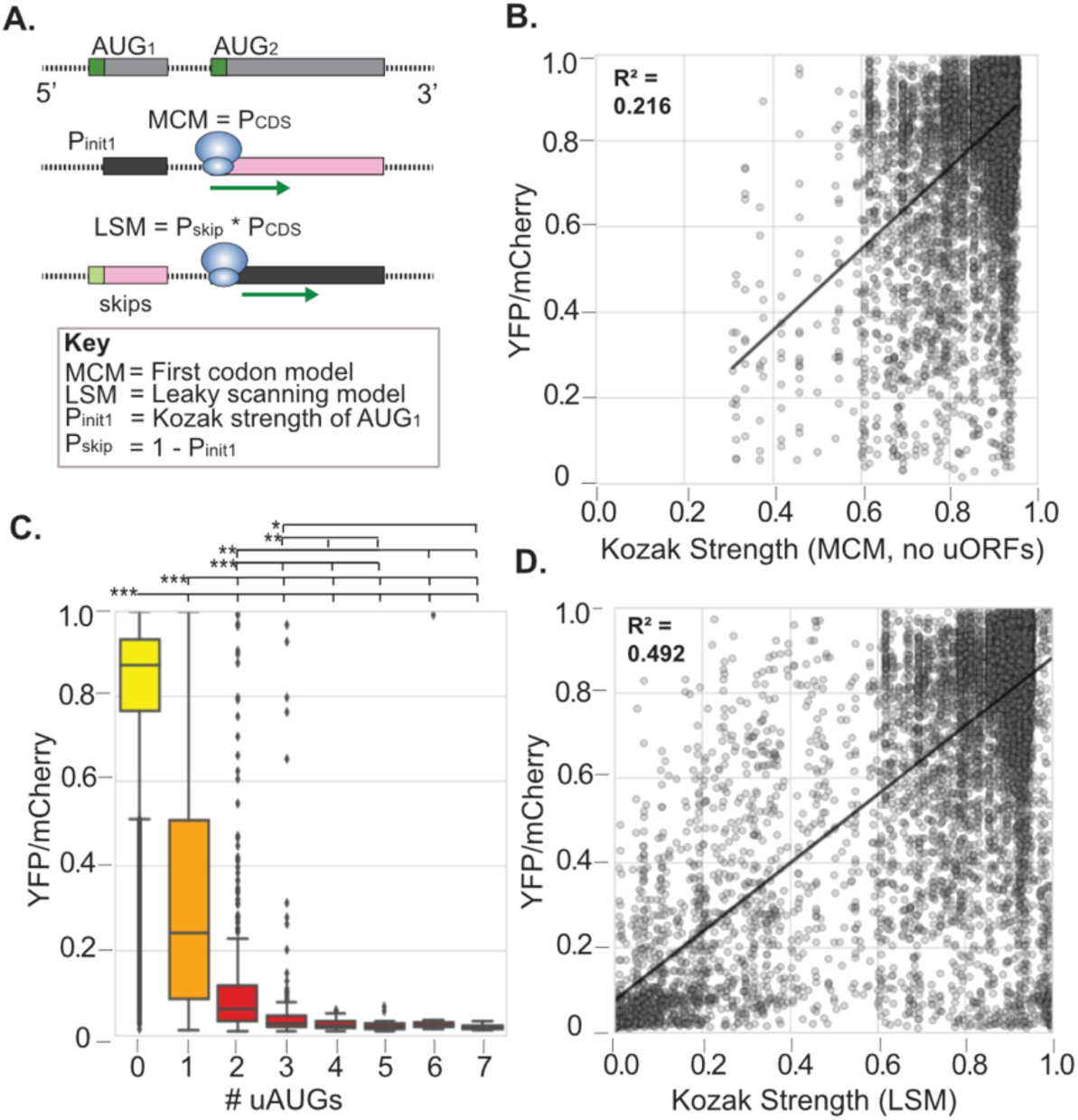
A simple leaky scanning model explains half of the variance of gene expression from natural yeast TLs. (A) Schematic describing a simple leaky scanning model. (Top) mRNA containing two AUG start codons. (Middle) The Main Codon Model (MCM) predicts YFP expression using the Kozak strength for the main CDS start codon. (Bottom) The Leaky Scanning Model (LSM) predicts YFP expression using the Kozak strengths of all AUGs. This example shows the ribosome skipping the first AUG and initiating at the second ORF (black). The probability of initiating at YFP is given by the fraction of ribosomes that reach the CDS start codon (Pskip) times the Kozak strength at the YFP start codon (Pinit2). (B) The MCM explains 21.6% of the variance in YFP levels for TLs without any upstream AUGs (R^2^ = 0.216; see Fig SX for uORF TLs). (C) Boxplots representing the distribution of measured YFP expression (y-axis) for native 5’ TLs binned by the number of uORFs (x-axis). Additional uAUGs further repress YFP expression. (P-values) *** < 0.001, ** < 0.01, * < 0.05. uAUGs:n = 0:9058, 1:1448, 2:309, 3:126, 4:50, 5:22, 6:10, 7:4 (D) Linear regression model of measured YFP (y-axis) versus the LSM predicted Kozak strength (x-axis) for each 5’ TL.

However, this did not include the impact of upstream AUGs (uAUGs) and their Kozak sequences during PIC scanning. In fact, the presence of even a single uAUG out-of-frame with the main ORF, significantly decreased expression (FIG 4C). Although rare in our library, TLs containing strong in-frame AUGs led to increased YFP expression as seen by several outliers, consistent with ribosome leaky scanning past annotated start codons. To incorporate uAUGs in Kozak predictions, we generated a Leaky Scanning Model (LSM) (FIG 4A). The LSM calculated probabilities of PICs initiating at uAUGs versus the main AUG based on Kozak strengths. Thus, the LSM captured the propensity of strong uAUG Kozak sequences to deter PICs from the main start codon. Meanwhile, weak Kozak sequences would permit PICs to bypass uAUGs and initiate at main AUGs with higher probability. Compared to the FCM, the LSM more accurately explained reporter expression with an R^2^ = 0.49 (FIG 4D). These results show a LSM using Kozak context measurements is a better predictor of gene expression. However, the discrepancy between the predicted and measured expression suggests that additional sequence features contribute to transcript leader regulation of gene expression.

### 5’ TLs features and Modeling YFP

To quantitatively compare the impacts of 5’ TL sequence features on gene expression, we constructed a predictive machine learning model. Given the variation in sequence length and our library size, we employed elastic net (EN) regression to evaluate the role of individual features on gene expression. We evaluated numerous TL sequence features, including Kozak context, uAUGs, G-quartets, and other structural and nucleotide composition features (FIG 5C, TABLE S3). We also included all nucleotide 4-mers, other known RNA binding protein motifs (35), and motifs recently reported to affect ribosome loading in *in vitro* extracts (12), resulting in over 200 features (TABLE S3). The EN selected and weighted all sequence features that contributed to a linear regression model of gene expression, together explaining 70% of the variance in gene expression (FIG 5A-B). We also constructed a model using more stringent parameters to identify robust features (Figure S3; R^2^=0.66). The most influential predictors of expression for our constructs, both under stringent and less stringent model conditions (see FIG 5B, FIG S4, and Table S4, Table S5), were Kozak strength (coefficient: 0.499, 0.542), presence of uAUGs (coefficient: -0.744, -0.486), and RNA levels (coefficient: 0.184, 0.458).

**Figure 5.**
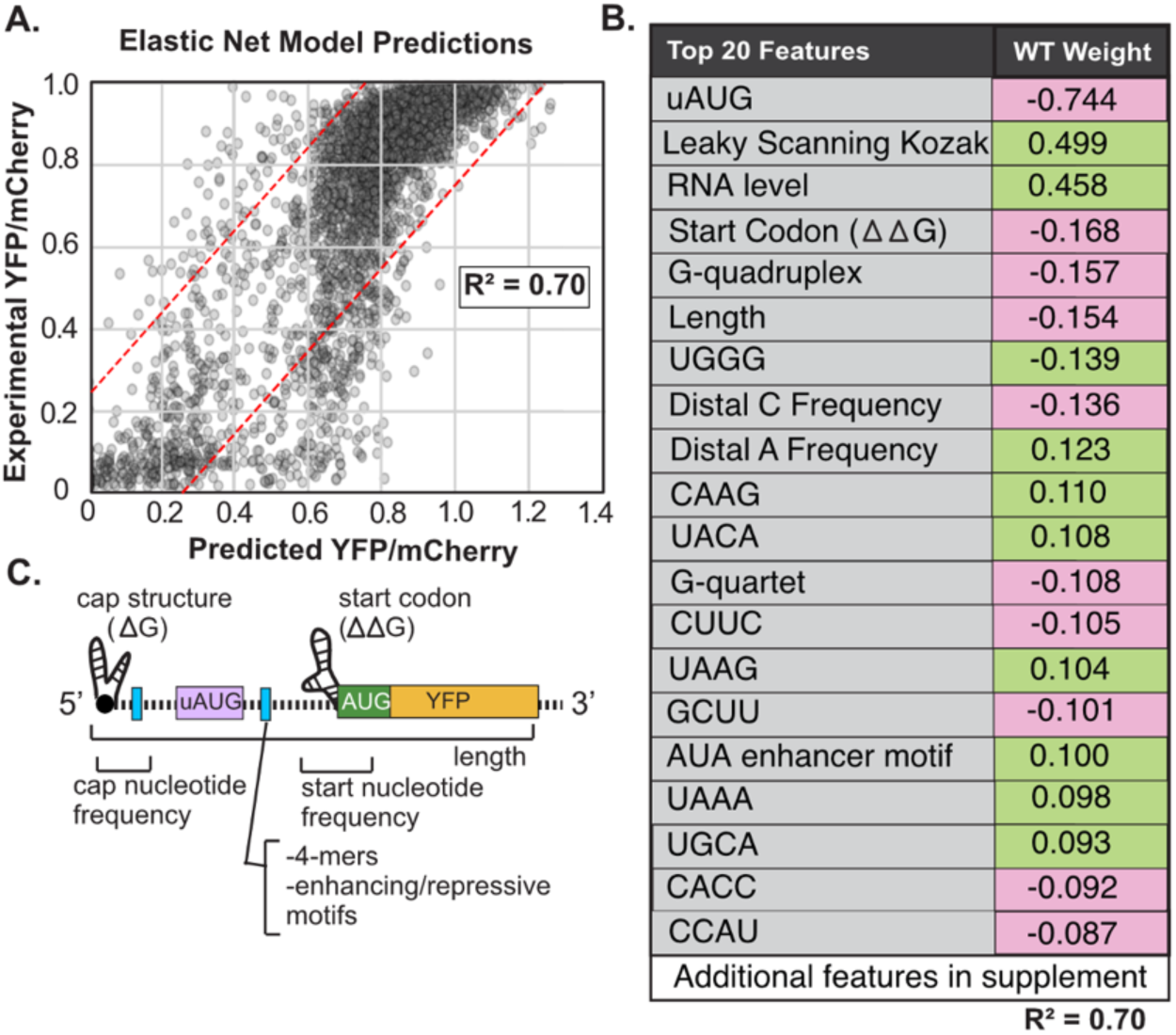
Elastic net regression determines the relative influence of transcript leader features on gene expression in vivo. (A) Elastic Net Model (EN) for predicting YFP expression. The scatter plot shows the measured YFP expression (y-axis) for all 5’ TLs versus the EN model predictions of YFP (x-axis) in WT yeast. The resulting model explains ∼70% of variance in experimental YFP. When the predictions are capped at 1, the resulting R^2^ drops to ∼0.69 (B) The table shows model coefficients for the significant features extracted from the EN model after n=100 iterations (additional features shown in supplemental data). (C) Schematic of 5’ TL features predicted to influence YFP expression based on the EN model.

The model also quantified the effects of mRNA folding around the 5’ cap and other 5’ TL on gene expression. For instance, structured 5’ caps are likely to hinder the binding of initiation factors, resulting in lowered PIC loading rates and decreased expression (FIG 5C) (1, 4, 43, 47). Additionally, strong structures around start codons could obstruct readability of the mainORF by impeding PIC scanning and efficient translation initiation (2, 4, 28). Moreover, the preference for increased adenine frequency (coef=0.123) around the start codon may aid in preventing the formation of strong structures. Several new 4mer motifs were identified *in vivo*, although their exact function is unknown. The G-rich 4mer sequences (e.g. TGGG) typically repressed YFP expression while AT-rich were positively correlated with expression (e.g. TAAA) (FIG 5B). From the EN analysis of our TL library, gene expression is usually negatively impacted by TL length. While the length of the TL sequence is independently associated with variance in gene expression, it also demonstrates a correlation with increased overall TL structure and other regulatory elements.

Recently, Niederer et al. investigated mechanisms driving ribosome recruitment to 5’ TLs lacking uAUGs *in vitro* using another library approach (DART) (12). By comparing the efficiency of ribosome recruitment to 5’ UTRs in translation extracts, they uncovered several enhancer and repressive motifs. Applying our EN model, we found that the AUA motif, previously reported as an enhancer by Niederer et al., significantly contributed (coef=∼0.1) to the variance in YFP expression, indicating a positive influence albeit with a somewhat modest effect size (FIG 5B). Our results were also consistent with other motifs from the previous *in vitro* study (12), although their relative impacts on gene expression *in vivo* were weaker, as indicated by their lower model weights. To more directly compare our results to Niederer et al., we reevaluated the EN model using TLs lacking uAUGs (FIG S5, R^2^=0.49; Table S6). The results reinforced the small positive impact (coef=0.063) that the AUA motif plays on expression, although the precise regulatory mechanism is unknown. These results show that motifs previously found to influence ribosome loading *in vitro* are associated with similar effects on gene expression *in vivo*, although they account for a modest amount of the variance among natural yeast transcript leaders.

We next examined sequence motifs in the EN outlier predictions (over-predicted or under-predicted) to improve our model. Outliers were defined as having a change of +-0.25 in measured versus predicted YFP expression (FIG 5A, dashed lines, methods). Our analysis via STREME (48) identified several A/U-rich motifs that were overrepresented in the outliers (FIG S6). Thus, these motifs may play roles in regulating transcription or translation. Despite incorporating these motifs into the model, we did not observe any increase in model performance, and the coefficients for the added motifs were not significant. These findings suggest that there may be unidentified sequence elements in TLs that impact expression levels. Overall, our EN models helped us quantify the relative impacts of *cis*-acting sequences and structures on expression from native yeast TLs *in vivo*.

### Impacts of alternative TSSs on protein levels

In eukaryotes, alternative TSSs can change gene expression levels by introducing additional sequence features. To investigate the regulatory capacity of yeast alternative TLs on gene expression, we compared YFP levels from alternative TSSs in the 5’ TL library. We calculated the differences in YFP expression for all pairwise alternative TSSs. We found that alternative TL isoforms influenced expression up to 78-fold (FIG 6A). Indeed, changes of as little as 15 nucleotides altered protein expression as much as ∼16 fold. Longer TL isoforms (∼2000 TLs) were generally less efficient at translation. In many cases, the longer TLs contain additional uAUGs or more stable RNA structures. For example, a 50 nucleotide longer isoform of the transcript leader from *ARG8*, an aminotransferase encoding gene, introduced a new uAUG with a strong -3A at the 5’ end (FIG 6B). Similarly, a ∼100 nt difference in *AST2* transcript leaders introduced four new uAUGs (FIG 6B). Three uAUGs were out-of-frame with the main start codon and thus competed for PIC recognition and initiation, while the other uAUG created an N-terminal extension. The longer *AST2* transcript is also predicted to form a G-quartet/G-quadruplex structure (46) which may further contribute to its repressive nature. Although rarer in our library, longer TLs sometimes drove increased YFP expression (26%: ∼600 TLs). The ubiquitin-specific protease, YER151C (UBP3) has two alternative transcription start sites that differ by 16 nt. While the sequence features are similar between the two transcripts, the longer isoform produces a less-structured cap which likely increases PIC 5’ end binding and scanning (43, 49). One outlier was YLR265C (*NEJ1*), where a 20nt longer TL increased expression 32-fold. The 5’ TL features did not change significantly; however, the distance from the cap to the first uORF increased, suggesting that uAUG location affects initiation as shown in May et al. (13). To further evaluate the accuracy of our EN model for transcript leaders, we compared the predicted and measured differences in expression from alternative TSSs. Notably, we found good agreement (R^2^ = 0.57; FIG 6C). Together, these results determine the breadth of expression differences caused by altering only the 5’ TSS in native yeast TLs *in vivo* and highlight TL features that could account for this variation.

**Figure 6.**
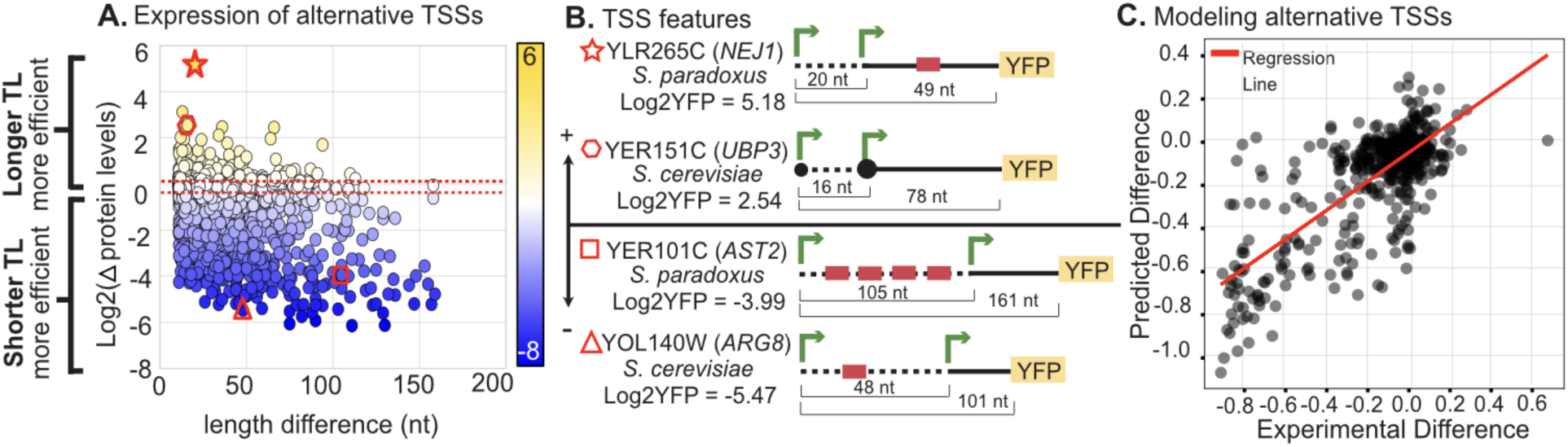
Effects of natural yeast alternative transcription start sites (aTSSs) on protein expression in vivo. (A) Scatter plot compares the log fold change (log2(Long/Short)) of 5’ TLs for genes with alternative TSSs (x-axis=length; y-axis=log2change; R^2^=0.153). Typically, longer TLs displayed greater changes in YFP levels. The negative log2YFP values (blue) indicate that the longer TL was less efficient at translation compared to its shorter counterpart. A positive log2YFP (yellow) values reveal that the shorter TL had lower YFP levels. The red dashed lines represent the median for positive (0.139) and negative (-0.411) values. Red symbols identify genes represented in FIG 2B. (B) Examples of alternative transcript leaders that significantly changed YFP expression. (B) YLR265C (S. paradoxus) and YER151C (S. cerevisiae) both had longer transcripts that increased YFP levels. (C) YOL140W (S. cerevisiae) and YER101C (S. paradoxus) were two examples of longer transcript leaders which repressed YFP expression with additions of uAUGs. R^2^ = 0.57.

## DISCUSSION

Due to their substantial influence on mRNA translation and decay, 5’ transcript leaders play a key role in regulating gene expression. Early studies mainly examined the effects of individual 5’ TL sequence elements, such as Kozak context and RNA structure, on gene expression (2, 43, 49, 50). More recently, massively parallel reporter assays (MPRAs) have allowed researchers to simultaneously test thousands of designer and randomized sequences *in vivo* (10, 14, 51–55) and *in vitro* (12). Although informative, these studies often involved technological compromises that could affect their interpretation. For instance, several studies appended long fixed sequences to the 5’ terminus of reporter libraries. While this facilitates PCR amplification, it also removes all variation in sequence and structure at the 5’ cap. Several previous studies also used fixed length, randomized TL sequences (10, 14, 51, 55). Consequently, such studies cannot capture the effects of TL length and have limited structural variation. Finally, previous large-scale studies of TL effects were limited to either protein expression or ribosome loading alone. Here, we combined three separate MPRA analyses to investigate the impact of 5’ UTR sequences on mRNA levels, ribosome recruitment, and protein expression. By assaying thousands of natural yeast transcript leaders, we quantified the respective roles of known and novel 5’ UTR *cis*-acting elements in gene expression.

The Kozak sequence context surrounding mRNA start codons has long been recognized as a major determinant of initiation efficiency (56). The strength of Kozak contexts influences translation initiation via leaky scanning, in which PICs bypass inefficient start codons in a condition-specific manner (41, 57). We evaluated a simple leaky-scanning model (LSM) incorporating all uAUGs and their Kozak contexts in our library of reporter elements. Our LSM explains ∼49% of variance in expression. This indicates that the Kozak sequence, though undoubtedly important, cannot fully explain the variation in translation initiation found in yeast 5’ UTRs. Notably, it has been suggested that the Kozak sequence is less impactful in *S. cerevisiae* than in other eukaryotes (58), due to loss of protein interaction domains in yeast initiation factors. Thus, leaky scanning may play a more prominent role in other species. Similarly, previous work found that near-AUG start codons can drive significant translation initiation in mammalian tissue culture cells (59), but not in yeast (34). Future work is needed to determine whether Kozak sequences, near-AUG start codons, and leaky scanning explain more of the variance in expression across natural human 5’ TLs than we observed with yeast TLs.

Although our LSM was a better predictor of yeast TL activity than start codon Kozak strength alone, 5’ TLs contain many *cis*-regulatory elements that could affect scanning PICs (57, 60). For example, including constraints on uORF start codon structure was found to greatly improve translation efficiency predictions for human SERPINA1 mRNA isoform reporters (28). Incorporation of isoform-specific structure probing data might increase the predictive power of our LSM. Indeed, we found several TL structural features influenced protein expression, including structure around the main ORF start codon and G-quadruplexes. In addition, many other features of uORFs can affect their relative influence on leaky-scanning, including uORF length, position, and the charge of encoded uORF peptides (34). By increasing the time ribosomes occupy 5’ TLs, these features may inhibit the loading and scanning by additional PICs. For example, PIC collisions or pausing may lead to slowed main ORF initiation, a shift of translation to upstream start codons, or increased mRNA decay (61–64). Such a reduction would give uORFs a greater influence on initiation than expected from leaky scanning models.

The limitations of LSM models may be addressable using Totally Asymmetric Simple Exclusion Process (TASEP) models (65, 66). For example, Andreev et al. described a TASEP model variant (ICIER) in which ribosomes translating uORFs displace downstream scanning PICs, while upstream scanning PICs queue behind translating ribosomes. Notably, this model can create automatic derepression of translation under stress conditions, when PICs load less frequently at mRNA 5’ ends. However, the model seems incompatible with studies that suggest mRNA 5’ ends remain tethered to scanning PICs (67–69), as this would not allow new PICs to load upstream of scanning PICs. Furthermore, the rapid nature of translation initiation (70) may make the presence of multiple PICs on individual TLs rare. Yet, TASEP-inspired models hold promise for future prediction of translation initiation, especially with the incorporation of additional parameters, including mRNA structures.

Unlike mechanistic models such as LSM and TASEP (71), computational and machine learning methods can identify more complex TL features and interactions (10, 11, 14, 72). Although previous studies investigated 5’ TL control and identified regulatory elements (10–12), they did not directly explain what occurs in native transcript leaders *in vivo*. To better explain natural gene expression, we built a comprehensive 5’ TL model by combining FACS-Seq with elastic net regression. The resulting model explained ∼70% of variation in YFP for all native yeast TLs up to 180 nucleotides *in vivo.* Our results confirmed known *cis*-acting motifs, including AUG-Kozak pairs, sequence composition, and mRNA structures, while also extending our understanding by defining their relative roles in natural TLs. By incorporating motifs previously reported to affect ribosome loading *in vitro* (12), we were able to confirm the *in vitro* “AUA” enhancer element *in vivo*. Several 4-mers were identified which may be indicative of structure or include un-identified RNA binding motifs. While it is unclear which *trans*-acting factors recognize the AUA containing motif, notably the motif is similar to the position specific element used in mRNA cleavage and 3’ end formation (73). We also identified C/U rich motifs associated with lower expression and ribosome loading. This motif is found in sequences bound by the translational repressor *SSD1* (*74*), making it a prime candidate for the corresponding *trans-*acting factor. Our model underscores the importance of the nucleotide composition at the 5’ cap and around the start codon which could influence structure and PIC binding and initiation. Thus, our EN models demonstrate the wide regulatory potential of 5’ TLs and their direct impacts on gene expression.

Although our EN model accurately predicts expression, it has some limitations. Our structural predictions are constrained as they rely on predictive ΔG measurements that fail to capture various mRNA structural states. To overcome this limitation in future studies, it would be valuable to verify mRNA structures and cap sequences via structure probing experiments. In a complementary paper, we discuss how uORF features, such as location, codon makeup, and length, impact expression (13). These features may account for some of the unexplained variance in our current LSM and EN models.

Furthermore, the hidden interactions between structures and uORFs are not directly examined. There may be a seesaw effect on the level of repression between these two features. Upstream structures can mask the repressive nature of uORFs by reducing PIC loading, occluding uORF starts, and minimizing PIC collisions or queuing on uORFs (29, 75). Indeed, we see some of the overpredicted TLs containing strong 5’ cap structures along with uORFs downstream. Conversely, strong uORFs without any inhibitory structures upstream may dominate PIC usage, resulting in lowered mainORF expression. Some studies hint at such models by examining ribosome loading on uORFs versus CDSs (76, 77). Nevertheless, PIC scanning and pausing are still not well understood. By studying PIC interactions or disomes, we may increase our understanding of scanning efficiency. Additionally, mRNAs interact with *trans-*acting factors such as 5’ RNA binding proteins, represented by consensus binding motifs in our model. Capturing RBP interactions through CLIP protocols could boost the predictability of our 5’ TL regulatory models. There may be other unknown features or feature interactions that explain the remaining 1/3 of variance in expression for our model.

Notably, we found a complex relationship between mRNA levels, translation, and protein expression. In general, TLs containing uORFs were associated with lower mRNA abundance and correspondingly lower protein levels. Consistent with this, we found TLs that had very low ribosome loading also had correspondingly low mRNA levels. However, we also observed low mRNA levels for TLs with the highest ribosome load. This suggests a model in which overloading of ribosomes leads to mRNA destabilization, perhaps via ribosome collisions and the ribosome quality control decay pathway (78, 79). In this case, TLs may be somewhat optimized for specific protein coding genes such that collision prone transcripts have reduced ribosome loading while rapidly translating ORFs can load ribosomes more quickly (80). If so, this would have important implications for the design of mRNA therapeutics, as the optimal rate of ribosome loading may depend on the probability of ribosome collisions in a given ORF.

Many of the alternative TSSs we tested are differentially regulated in yeast. Previous work suggests such transcripts may have regulatory roles. For example, shifts in TSS usage have been reported in different environmental conditions (30) and throughout meiosis (16, 81). Studies in mammals suggest that alternative TSSs drive mRNA isoform diversity and have specific functions (82–84). Our analyses showed that yeast alternative transcript leaders can alter protein expression up to ∼80-fold, revealing the vast regulatory potential and diversity of yeast 5’ TLs. Although upstream TSSs generally produced less efficient TLs, there were cases where we observed increases in expression.

Further, the changes in alternative TSS expression were frequently attributed to the introduction or removal of regulatory features. Future work is needed to evaluate the production and significance of alternative TSSs. Such studies are expected to reveal regulatory links between transcription regulation, environmental conditions, mRNA translation and turnover.

In summary, we evaluated the regulatory roles of TL features in yeast grown to log phase under unstressed conditions. However, environmental stimuli can cause dramatic changes in mRNA translation, including suppressing canonical cap translation and altering the effects of sequence features (85–90). While current experiments of single genes under stress or varied conditions provide valuable insights, large-scale studies are needed to detect critical TL features and their role in genetic reprogramming. New methods, such as incorporation of rapidly turned over proteins or degrons (91–93) may be useful in stress response studies. It would also be interesting to investigate the direct impact of stress and the interplay of TL features on PIC scanning. Such future studies of translational response to stress may provide key-insights into reestablishing homeostasis, channeling resources towards stress-related genes, and altering PIC trajectories.

## DATA AVAILABILITY

The raw sequencing data for FACS-uORF, PoLib-seq, and mRNA levels were downloaded from NCBI under the SRA accession number PRJNA721222(34)

## SUPPLEMENTARY DATA

Supplementary figures are shown below. Supplementary data tables will be available online at the journal website.

## ACKNOWLEDGEMENTS

The authors would like to thank the members of the McManus and Woolford laboratories at CMU for helpful discussions and suggestions. This study was supported by NIGMS grant R35GM145317 to CJM.

---

**FIGURE S1.**
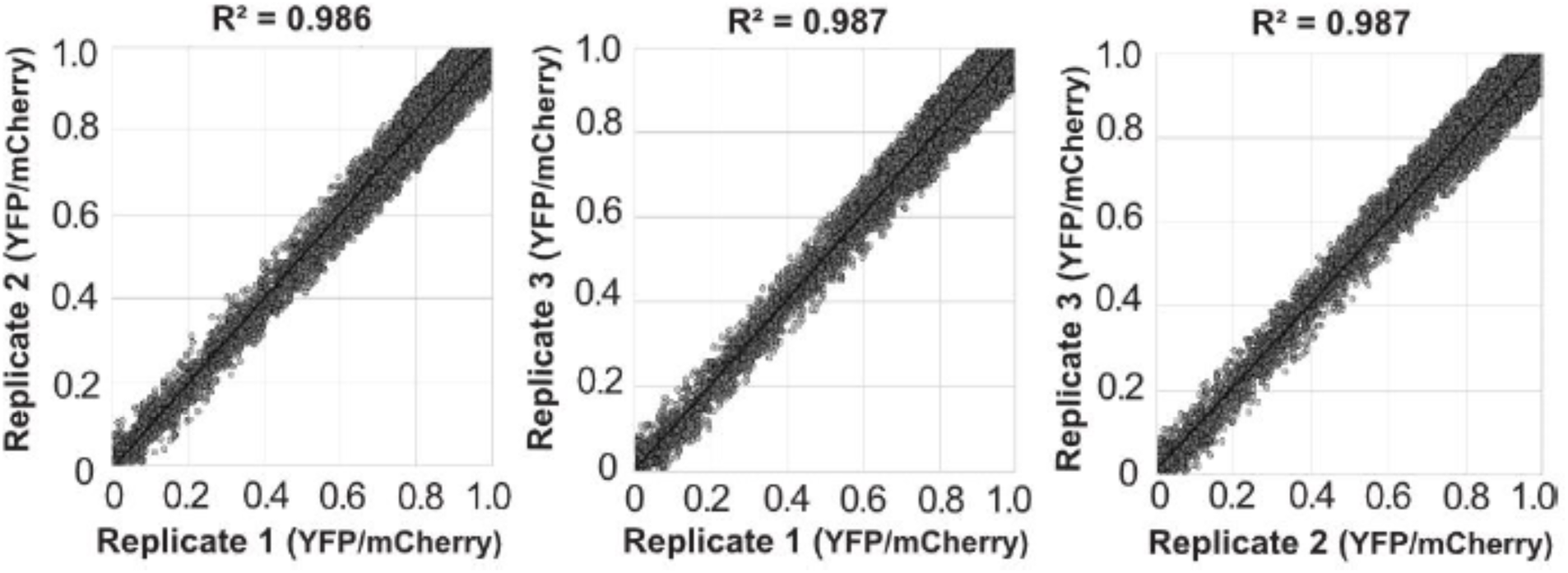
Reproducibility of FACS-Seq measurements of mean YFP levels for 5’ TLs. The scatter plots show the comparison of three replicates. R^2^ values are listed above each plot. Note that reporters with noisy expression measurements (coefficient of variation > 0.05) were removed from this and other analyses in this study, as previously reported (34).

**FIGURE S2.**
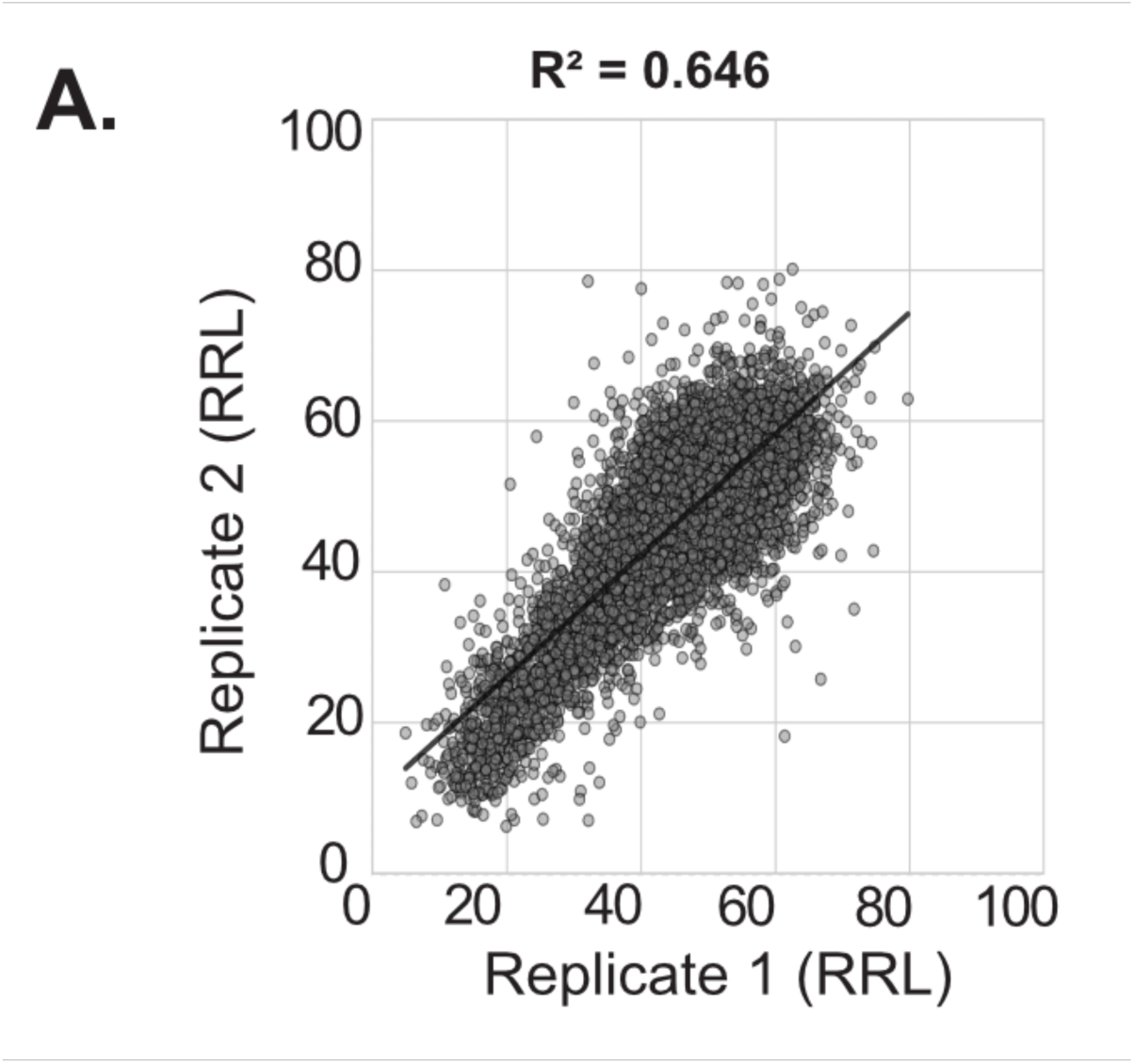
Reproducibility of PoLib-Seq measurements of ribosome loading levels for 5’ TLs. The scatter plots show the comparison of two replicates with the R^2^ value listed above the plot.

**FIGURE S3.**
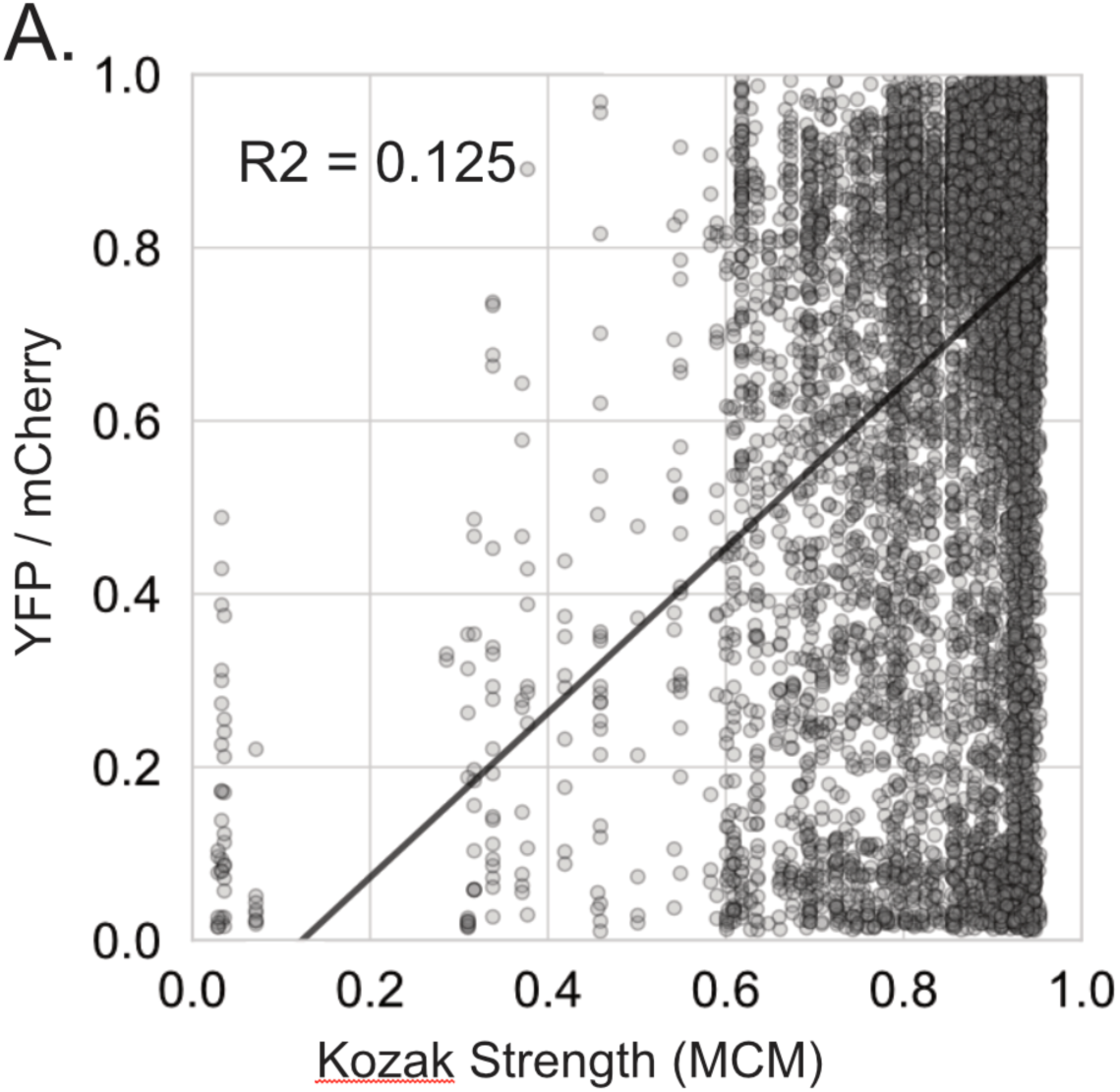
The Main Codon Model (MCM) Kozak strength performs poorly at predicting YFP levels across yeast 5’ TLs. This analysis included all TLs, including those with uAUGs and uORFs.

**FIGURE S4.**
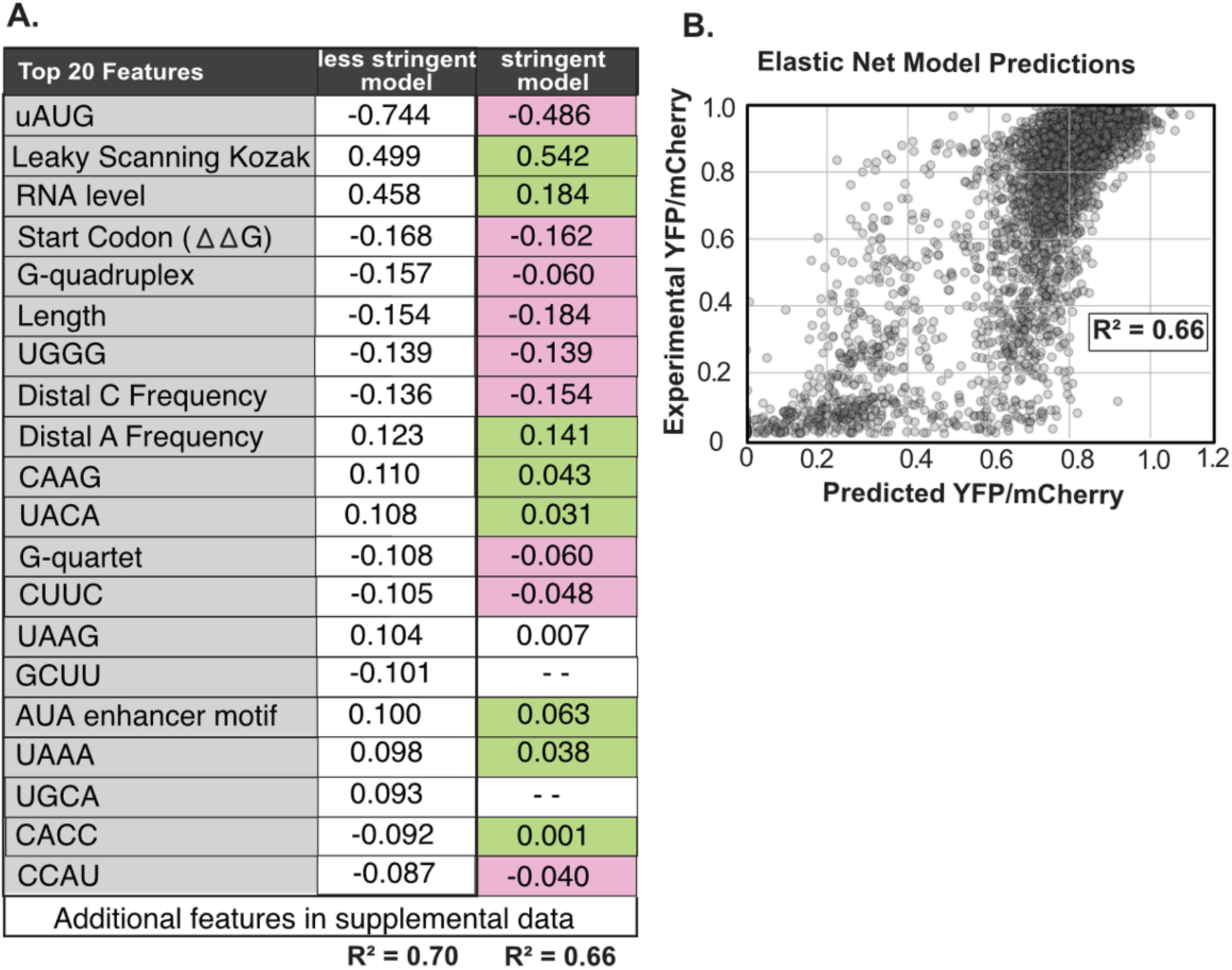
Comparison of less stringent and stringent elastic net model (A) The table comparing model coefficients for the significant features extracted from the less stringent EN model versus the stringent EN model after n=100 iterations (additional features shown in TABLE S5). The stringent model is similar, although several features are no longer significant (B) Elastic Net Model (EN) for predicting YFP expression. The scatter plot shows the measured YFP expression (y-axis) for all 5’ TLs versus the elastic net model predictions of YFP (x-axis) in WT yeast. The resulting model explains ∼66% of variance in experimental YFP.

**FIGURE S5.**
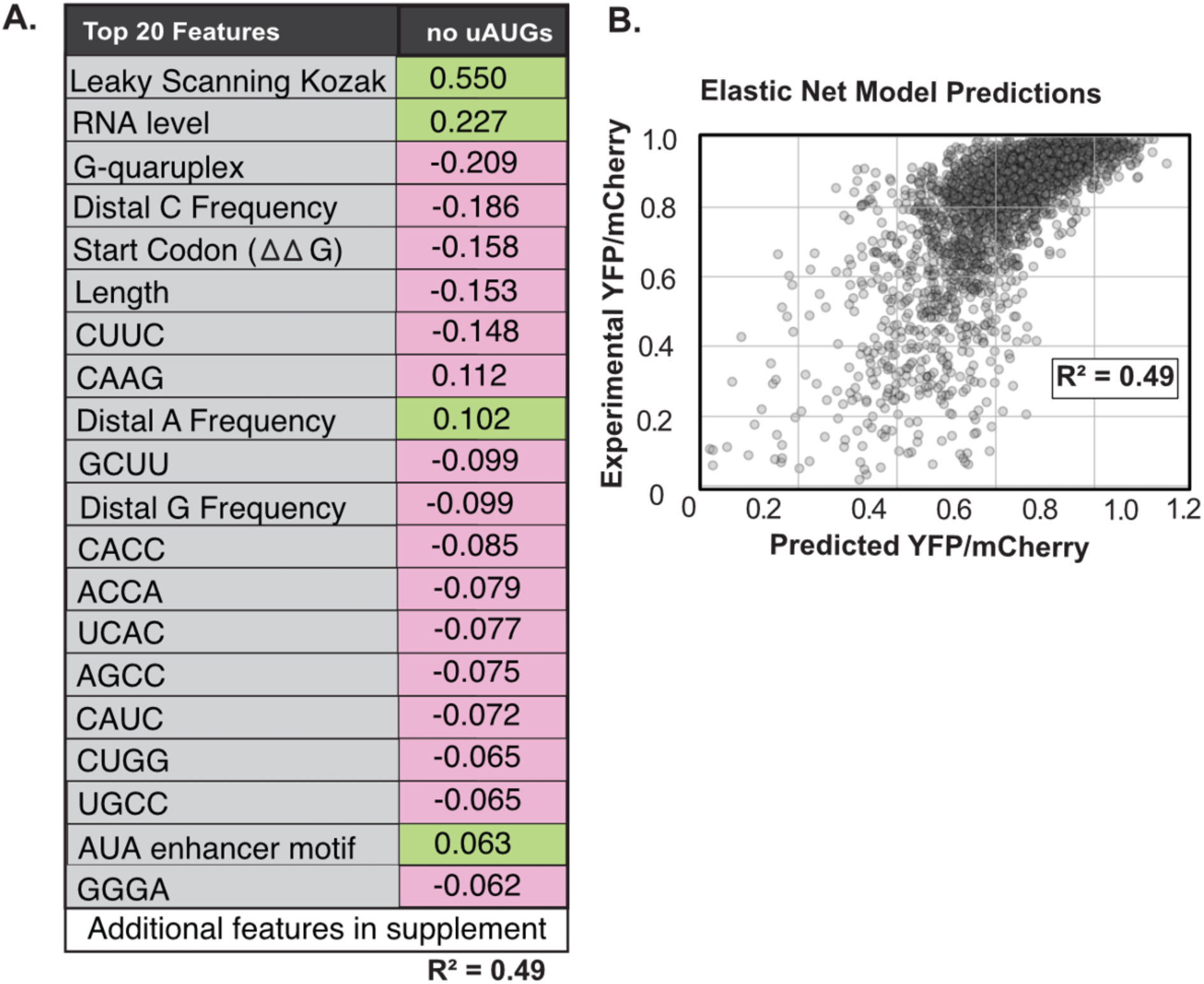
Elastic net model of TLs lacking uAUGs (A) Model coefficients for the significant features extracted from the less stringent EN model versus the stringent EN model after n=100 iterations (additional features x < 0.90 shown in TABLE S6). (B) Elastic Net Model (EN) for predicting YFP expression. The scatter plot shows the measured YFP expression (y-axis) for all 5’ TLs versus the elastic net model predictions of YFP (x-axis) in WT yeast. The resulting model explains ∼49% of variance in experimental YFP.

**FIGURE S6.**
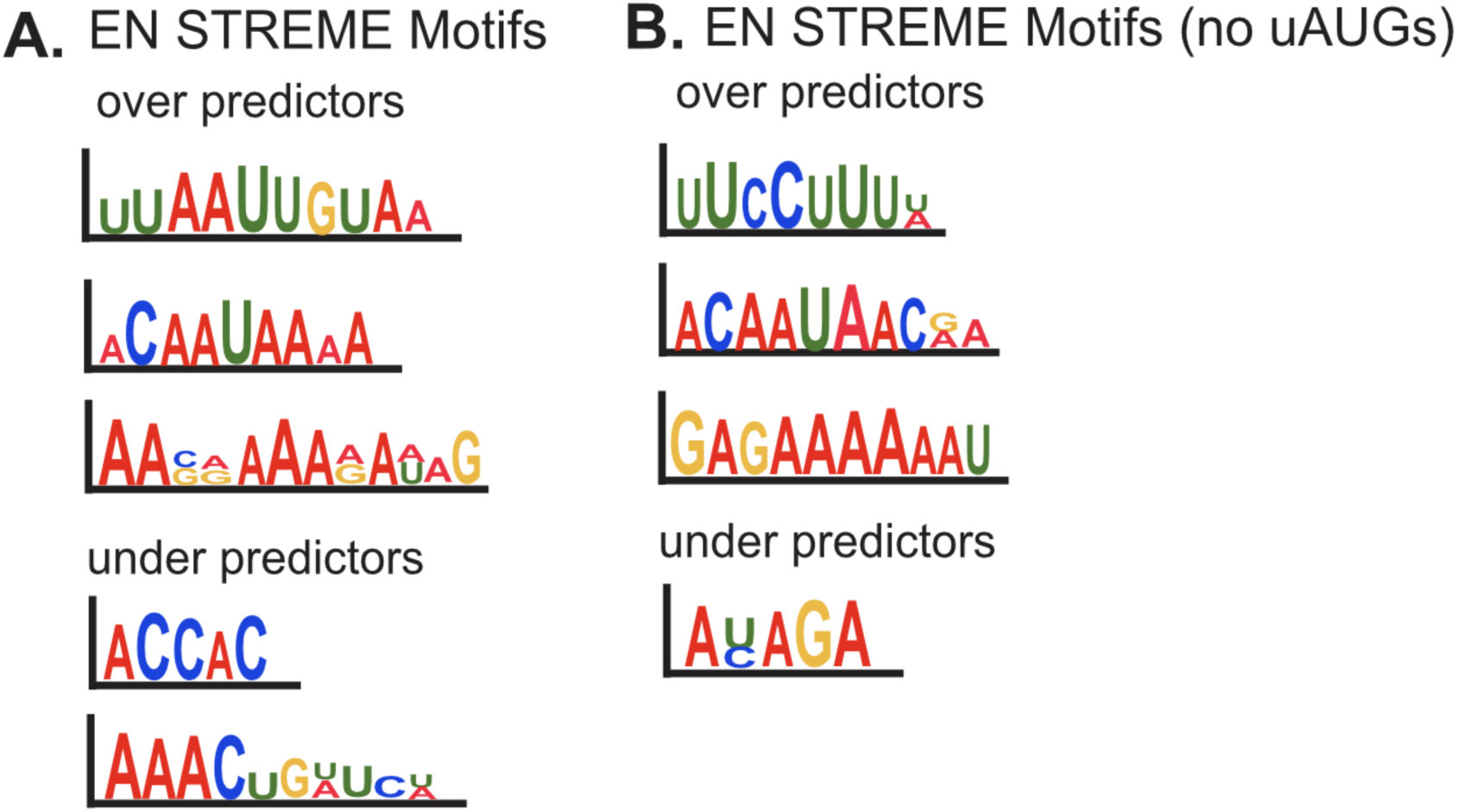
STREME motifs identified in the over- and under-predictors for the EN model with and without uAUGs. Over-predictors were labeled as +0.25 or above, while under-predictors were -0.25 or below. These outliers were calculated by measuring the change in measured vs. predicted YFP expression.

