## Supplementary Figures for "Deciphering the *cis-*regulatory landscape of natural yeast Transcript Leaders"

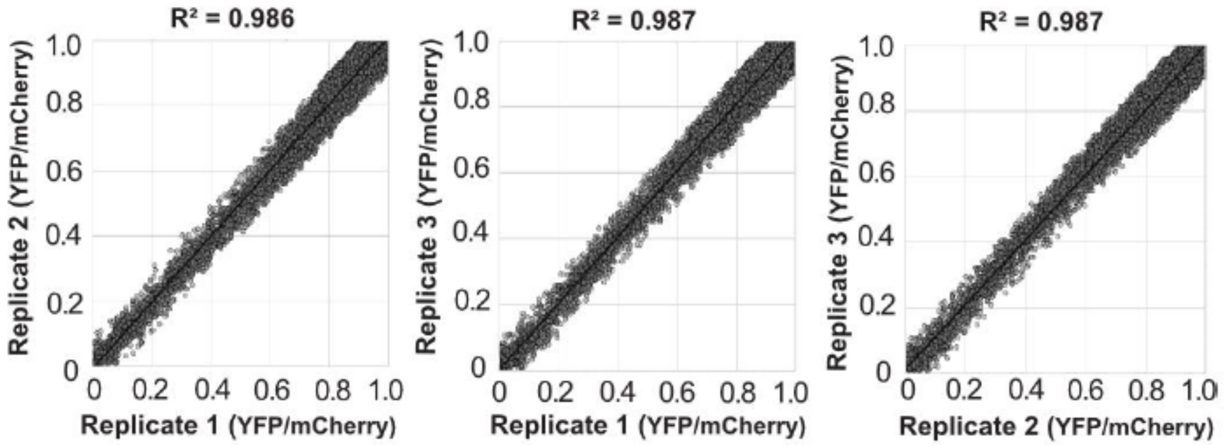


FIGURE S1 | Reproducibility of FACS-Seq measurements of mean YFP levels for 5′ TLs. The scatter plots show the comparison of three replicates. $R^2$ values are listed above each plot. Note that reporters with noisy expression measurements (coefficient of variation > 0.05) were removed from this and other analyses in this study, as previously reported (34).

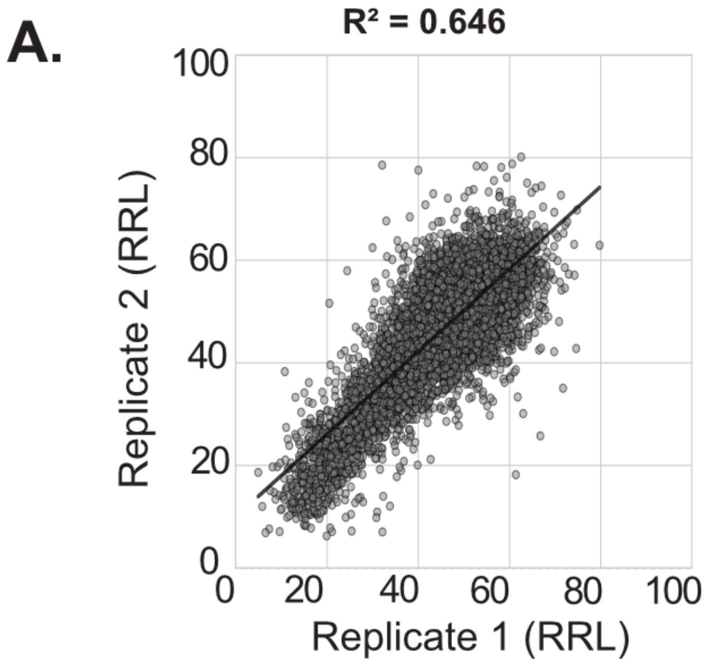


FIGURE S2 | Reproducibility of PoLib-Seq measurements of ribosome loading levels for 5’ TLs. The scatter plots show the comparison of two replicates with the $R^2$ value listed above the plot.

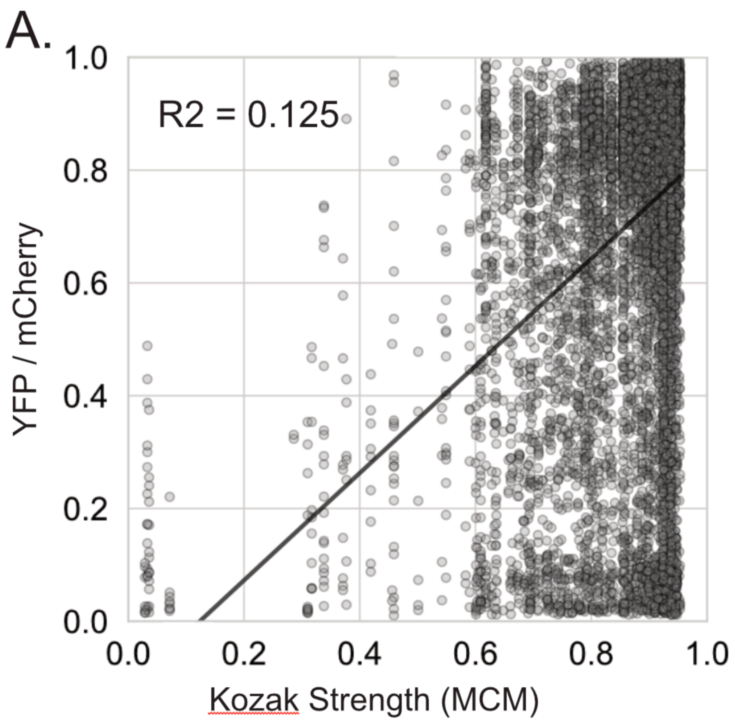


FIGURE S3 | The Main Codon Model (MCM) Kozak strength performs poorly at predicting YFP levels across yeast 5' TLs. This analysis included all TLs, including those with uAUGs and uORFs.

**A.**

| Top 20 Features | less stringent model | stringent model |
|---|---|---|
| uAUG | -0.744 | -0.486 |
| Leaky Scanning Kozak | 0.499 | 0.542 |
| RNA level | 0.458 | 0.184 |
| Start Codon ($\Delta\Delta G$) | -0.168 | -0.162 |
| G-quadruplex | -0.157 | -0.060 |
| Length | -0.154 | -0.184 |
| UGGG | -0.139 | -0.139 |
| Distal C Frequency | -0.136 | -0.154 |
| Distal A Frequency | 0.123 | 0.141 |
| CAAG | 0.110 | 0.043 |
| UACA | 0.108 | 0.031 |
| G-quartet | -0.108 | -0.060 |
| CUUC | -0.105 | -0.048 |
| UAAG | 0.104 | 0.007 |
| GCUU | -0.101 | - - |
| AUA enhancer motif | 0.100 | 0.063 |
| UAAA | 0.098 | 0.038 |
| UGCA | 0.093 | - - |
| CACC | -0.092 | 0.001 |
| CCAU | -0.087 | -0.040 |
| Additional features in supplemental data | | |
| | $R^2 = 0.70$ | $R^2 = 0.66$ |

**B.**

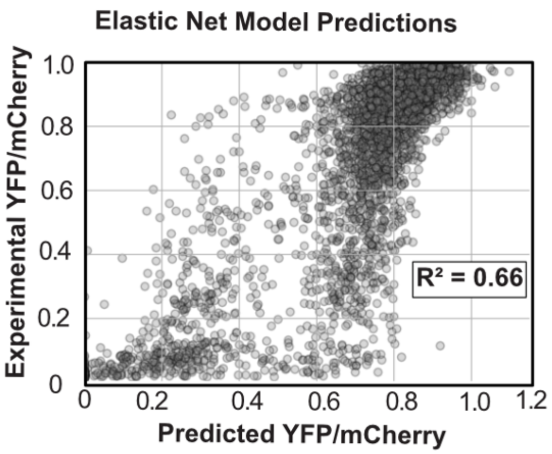


FIGURE S4 | Comparison of less stringent and stringent elastic net model (A) The table comparing model coefficients for the significant features extracted from the less stringent EN model versus the stringent EN model after n=100 iterations (additional features shown in TABLE S5). The stringent model is similar, although several features are no longer significant (B) Elastic Net Model (EN) for predicting YFP expression. The scatter plot shows the measured YFP expression (y-axis) for all 5' TLs versus the elastic net model predictions of YFP (x-axis) in WT yeast. The resulting model explains ~66% of variance in experimental YFP.

A.

| Top 20 Features | no uAUGs |
|---|---|
| Leaky Scanning Kozak | 0.550 |
| RNA level | 0.227 |
| G-quaruplex | -0.209 |
| Distal C Frequency | -0.186 |
| Start Codon (ΔΔ G) | -0.158 |
| Length | -0.153 |
| CUUC | -0.148 |
| CAAG | 0.112 |
| Distal A Frequency | 0.102 |
| GCUU | -0.099 |
| Distal G Frequency | -0.099 |
| CACC | -0.085 |
| ACCA | -0.079 |
| UCAC | -0.077 |
| AGCC | -0.075 |
| CAUC | -0.072 |
| CUGG | -0.065 |
| UGCC | -0.065 |
| AUA enhancer motif | 0.063 |
| GGGA | -0.062 |
| Additional features in supplement | |

$R^2 = 0.49$

B.

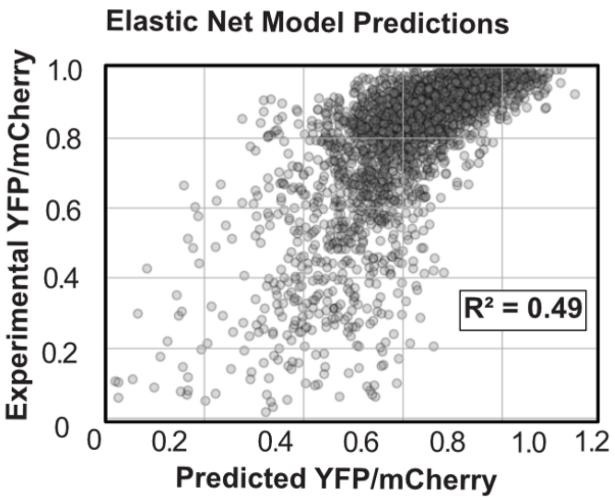


FIGURE S5 | Elastic net model of TLs lacking uAUGs (A) Model coefficients for the significant features extracted from the less stringent EN model versus the stringent EN model after n=100 iterations (additional features x < 0.90 shown in TABLE S6). (B) Elastic Net Model (EN) for predicting YFP expression. The scatter plot shows the measured YFP expression (y-axis) for all 5' TLs versus the elastic net model predictions of YFP (x-axis) in WT yeast. The resulting model explains ~49% of variance in experimental YFP.

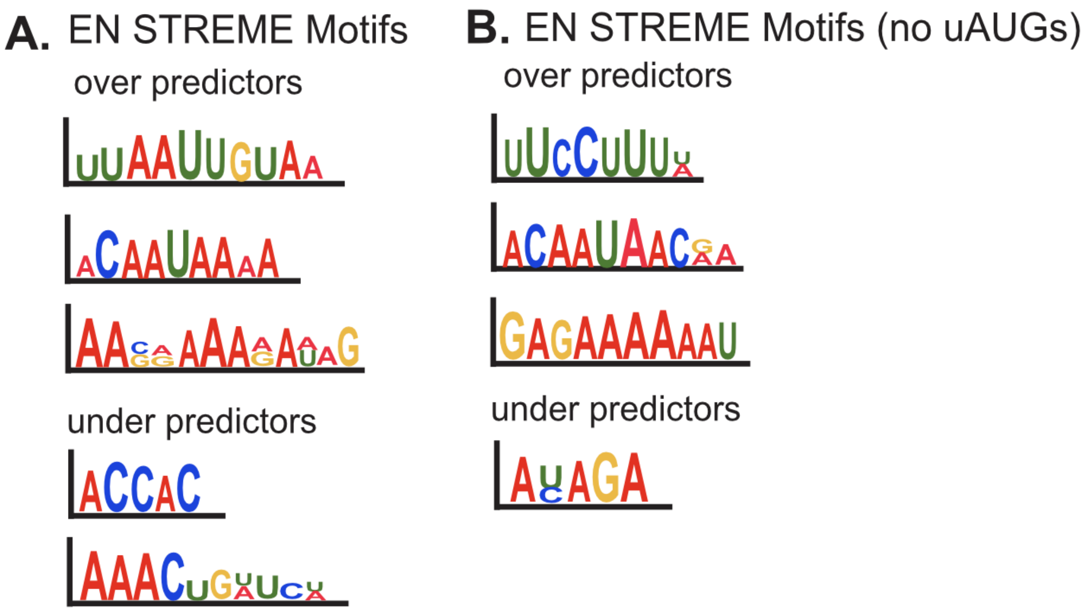


FIGURE S6 | STREME motifs identified in the over- and under-predictors for the EN model with and without uAUGs. Over-predictors were labeled as +0.25 or above, while under-predictors were -0.25 or below. These outliers were calculated by measuring the change in measured vs. predicted YFP expression.
